# Plumage redness signals mitochondrial function in the House Finch

**DOI:** 10.1101/728873

**Authors:** Geoffrey E. Hill, Wendy R. Hood, Zhiyuan Ge, Rhys Grinter, Chris Greening, James D. Johnson, Noel R. Park, Halie A. Taylor, Victoria A. Andreasen, Matthew J. Powers, Nicholas M. Justyn, Hailey A. Parry, Andreas N. Kavazis, Yufeng Zhang

## Abstract

Carotenoid coloration is widely recognized as a signal of individual condition in various animals, but despite decades of study, the mechanisms that link carotenoid coloration to condition remain unresolved. Most birds with red feathers convert yellow dietary carotenoids to red carotenoids in an oxidation process requiring the gene encoding the putative cytochrome P450 enzyme CYP2J19. Here, we tested the hypothesis that the process of carotenoid oxidation and feather pigmentation is functionally linked to mitochondrial performance. Consistent with this hypothesis, we observed high levels of red ketolated carotenoids associated with the hepatic mitochondria of molting wild house finches (*Haemorhous mexicanus*), and upon fractionation, we found the highest concentration of ketolated carotenoids in the inner mitochondrial membrane. We further found that the redness of growing feathers was positively related to the performance of liver mitochondria. Structural modeling of CYP2J19 supports a direct role of this protein in carotenoid ketolation that may be functionally linked to cellular respiration. These observations suggest that feather coloration serves as a signal of core functionality through inexorable links to cellular respiration in the mitochondria.

## 1. Introduction

Carotenoids are responsible for the bright red, orange, and yellow coloration of many animal species, and this coloration serves as an important social signal of individual condition [1,2]. In many vertebrate species, individuals that display red-shifted coloration gain a mating advantage or hold more resources [3,4], and carotenoid coloration is among the most commonly cited example of a condition-dependent sexual ornament [5–7]. Compared to animals with less red ornamentation, individuals with redder ornaments are better at resisting and recovering from parasites and managing oxidative stress, among other measures of performance [8–11].

Despite decades of study, however, the mechanisms that link red carotenoid pigmentation to individual performance remain uncertain [12–15]. Hypotheses for how carotenoid coloration serves as a signal of individual condition have traditionally focused on resource limitations or the need to trade carotenoid usage to support physiological functions over ornamentation [16,17], but empirical support for both of these ideas is equivocal [12,18]. More recently, it has been proposed that coloration is controlled by metabolic function and in turn mitochondrial efficiency [9,19]. In support of this idea, a recent meta-analysis revealed that the link between color expression and individual condition is strongest in bird species that rely on the metabolic conversion of carotenoids for color displays [7]. Such carotenoid conversions are most relevant to red color displays because most animals that display red carotenoid coloration ingest only yellow carotenoids that they oxidize to red pigments in a process that is hypothesized to take place in mitochondria [19,20]. In this regard, it was recently shown that genes encoding proteins from the cytochrome P450 monooxygenase superfamily are required for the production of red ornamental carotenoids in birds [21–23].

In this study, we explored the hypothesis that red carotenoid coloration serves as a signal of mitochondrial performance. To do so, we compared the relationships between carotenoid coloration and mitochondrial performance in male house finches (*Haemorhous mexicanus*) that were actively producing ornamental feather coloration (figure 1). To produce red feather coloration used to attract females, house finches oxidize the yellow dietary carotenoid cryptoxanthin to the red pigment 3-hydroxyechinenone (3EH) in a process requiring the gene encoding the cytochrome P450 monooxygenase CYP2J19 [21,24]. Our central hypothesis is that the efficiency of the oxidation of yellow dietary pigments, and hence the coloration of feathers, is controlled either directly or indirectly by mitochondrial function (figure 2) [19,25]. To test this hypothesis, we performed a fractionation study to confirm the localization of red carotenoids in hepatic mitochondria. We subsequently compared several measures of mitochondrial performance in hepatic tissue of wild birds with different hues. Finally, we used this information, in conjunction with a molecular model of CYP2J19, to propose a hypothesis for how mitochondrial function may control carotenoid conversion.

**Figure 1.**
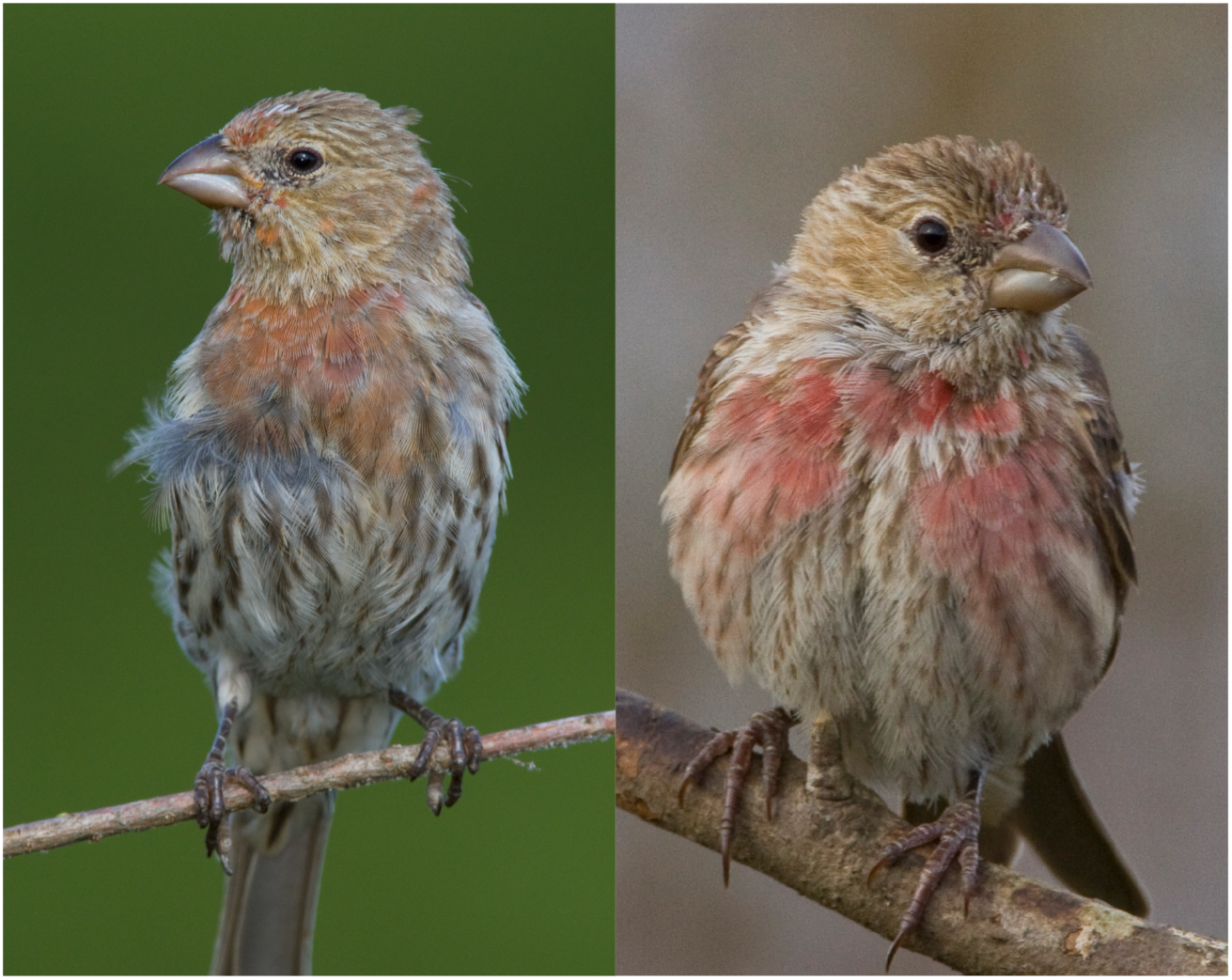
Hatching year house finches undergoing a first pre-basic molt, similar to those used in this study. The bird on the left is similar in hue to the drabbest male included in this study while the bird on the right is similar in hue to the brightest male included.

**Figure 2.**
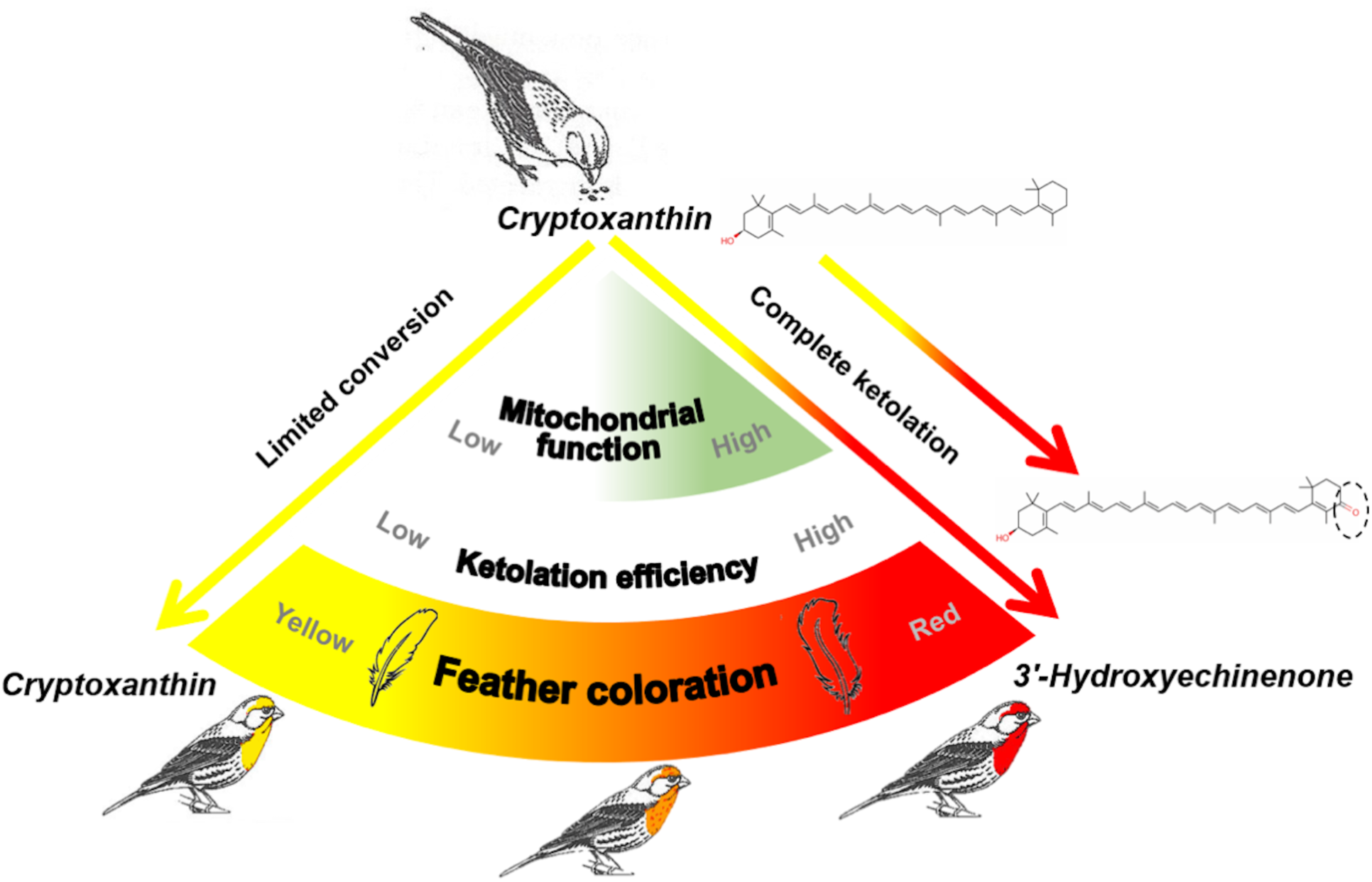
A schematic summary of the hypothesized links between red feather coloration and mitochondrial function. To produce red feathers, house finches ingest the yellow carotenoid cryptoxanthin and oxidize it to the red pigment 3-hydroxyechinenone. We hypothesized that ketolation efficiency is linked to mitochondrial bioenergetics. By this hypothesis, birds with low mitochondrial function have limited ketolation capacity and produce yellow feathers. Birds with high mitochondrial function have improved ketolation capacity and produce red feathers.

## 2. Materials and Methods

### (a) Field Collection

All procedures in this study were approved by the Auburn University Institutional Animal Care and Use Committee (PRN 2016-2922) and under federal (MB784373-0) and Alabama (6285940) collecting permits. We captured wild house finches at feeding stations using a walk-in basket trap as described in Hill [26]. All birds included in this study were males in the hatching year, transitioning from juvenal to first basic plumage via prebasic molt. In the juvenal plumage, house finches have no feathers with carotenoid pigmentation, so any red, orange, or yellow feathers in the breast plumage of birds had recently been grown. It was crucial to use birds in the process of molt and hence actively engaged in the production of red feather pigments, because this approach allowed us to match the current physiological state of birds to ornamentation that was actively being produced.

We captured birds at seven locations in Lee County, Alabama using large basket traps in which we suspended a tube feeder. Birds were captured between 07:00 and 10:00 on 11 mornings from July 20 to August 20, 2017 and 17 mornings from July 20 to August 17, 2018. Our protocol was to rapidly approach a trap holding birds—finches are undisturbed in these large traps until approached— and then to quickly move the birds from trap to brown paper bags. Paper bags enable birds to stand in a less stressful position and block all threatening visual and most auditory stimulation [26]. Thus, birds remained relatively calm between the time of capture and a maximum of three hours later when we removed them from the bag. Birds were taken from the bags, immediately anesthetized with isoflurane vapors and then sacrificed to collect tissues for physiological analyses.

### (b) Color analysis

The coloration of growing breast feathers was quantified from digital images of the ventral plumage of carcasses that were taken under standardized lighting with a color standard in each image. We quantified color from digital images instead of directly from feathers using a spectrometer because some birds used in this study had grown only scattered colored feathers. The human eye could see the color of incoming feathers and digital camera images captured the coloration, but there were no colored regions large enough to allow for accurate measurement with a spectrometer. Color quantification from digital images is reliable and repeatable [27,28].

We used Adobe Photoshop to first standardize all images to color standards within each image and then used the color sampler tool (Adobe Photoshop CS3 extended, v. 10.0, Adobe Systems, San Jose, CA, USA) to quantify the hue and saturation of feathers with carotenoid pigmentation at three points in each photo. Molting birds have uneven coloration and we focused color quantification on the largest unbroken patches of red/orange/yellow coloration where feathers were positioned to create a plane of coloration relative to the camera angle. Color patches were preselected before moving the selection wand over them and then the wand was moved to the center of the chosen patch and hue and saturation recorded. The PI who made the color measurements was not blind to the hypotheses being tested but was blind to the physiological measures for the birds being assessed; hence subconscious bias toward the hypothesis could not have affected the measurements of color. We averaged the three color measurements to arrive at a single hue value for each bird.

### (c) Tissue collection

A total of 91 birds were used in the study. We conducted fractionation of homogenized liver tissue using 55 of these birds. Fractionation of mitochondria required seven or eight times the tissue mass of the average mass of a House Finch liver. Thus, the livers of seven or eight males were pooled for each fractionation analyses. We pooled all hatching year, molting males captured in a particular day; hence, whether birds were added to a pool was not related to coloration or other variables. We used seven birds to create a single pool during the molt window of August 2017, and we used 48 birds each to create six pools, each composed of 8 birds, in the summer of 2018. We increased the number of individuals pooled between years because seven males used in 2017 provided barely enough tissue. A small cube of the right lobe of the liver was removed from 36 additional birds (collected in July-August 2017), flash frozen in liquid nitrogen, and then stored at -80°C for later analyses. The left and remaining right lobe of the liver from these birds was used for mitochondrial isolation and immediate testing of mitochondrial respiration.

### (d) Fractionation of mitochondria

Within 20 s of dissection from carcasses, livers were rinsed and minced together in SMEE (10 mM sucrose, 250 mM MOPS, 1 mM EDTA, and 1 mM EGTA) isolation buffer. During the entire procedure, all liver mitochondria preparations were kept at 4°C. Minced livers were then homogenized with a 2 mL glass Teflon homogenizer at 300 mg tissue per milliliter of buffer. Following the protocol of Trounce et al [29], as modified by Ingraham et al [29], the homogenate was centrifuged at 750 *g* for 10 min. The supernatant was then centrifuged at 9,800 *g* for 15 min. The resulting supernatant was saved for further separation into cytosol and microsome fractions (labeled ER-1). The microsome fraction was comprised primarily of fragments of endoplasmic reticulum (ER). The pellet was resuspended slowly by drop-wise addition of SMEE. This suspension was then centrifuged again at 9,800 *g* for 10 min. The pellet was resuspended in SMEE buffer and layered on top of 30% Percoll SMEE solution and subjected to ultracentrifugation at 150,000 *g* for 60 min along with the supernatant ER-1 from the 9,800 *g* spin containing cytosol and microsomes. The mitochondrial-rich layer from the Percoll solution was carefully harvested and washed 3 times with centrifugation at 7000 *g* for 10 min in SMEE at 4°C. The supernatant from ER-1 was saved as cytosol, and the pellet as microsome fraction.

For sub-mitochondrial fractionation, we followed a method previously described by Palczewski et al [30], with slight modification. Isolated mitochondria were diluted with SMEE to a minimum concentration of 50 mg mitochondrial protein/ml. Digitonin was added to a final concentration of 0.12 mg/mg protein. The solution was stirred on ice for 2 min and then diluted with 1.5 volume of SMEE. The solution was then centrifuged at 12,000 *g* for 10 min. The supernatant, which contained the outer mitochondrial membrane, was saved. The pellet, which contained mitoplast fraction, was resuspended in SMEE and then sonicated in an ice bath for 30 s (4 s on, 10 s off cycles). The sonicated material, as well as the previously saved supernatant, were then ultracentrifuged at 150,000 *g* for 60 min at 4°C.

We predicted that the ultracentrifuged pellet from the mitoplast fraction contained the inner mitochondrial membrane, while the supernatant from the same fraction contained the matrix. With this method, however, instead of having only inner mitochondrial membrane, the pellet should be considered a mixture of mostly inner mitochondrial membrane and some matrix content. The pellet from digitonin-treated supernatant contained outer mitochondrial membrane. Using the cubes of the liver frozen previously, fractions were then verified by immunoblot against subunits of carnitine palmitoyltransferase 1A (CPT1A), citrate synthase (CS) and cytochrome *c* oxidase (COX IV). In addition to detecting ketolated carotenoids in the inner mitochondrial membrane (IMM) and to lesser extent, the outer mitochondrial membrane (OMM), we also detected ketolated carotenoids in layer ER-1 (= 3.41+1.38 SD µg mL-1 3HE). This fraction is predicted to represent the endoplasmic reticulum-rich microsome. We did not confirm the identity of this layer with a protein marker so we cannot be confident in its identity. Observing ketolated carotenoids in a microsome layer would be expected if carotenoids are ketolated in or transported to the IMM because once ketolated, the pigments must be packaged for transport out of hepatic cells following ketolation.

### (e) Mitochondria measurements

Mitochondria were isolated following procedures outlined previously [31]. The fresh liver was minced and then homogenized in a Potter-Elvhjem PTFE pestle and glass tube. The resulting homogenate was centrifuged at 500 *g* for 10 minutes and the supernatant was then decanted through cheesecloth and centrifuged at 3,500 *g* for 10 minutes. The resulting supernatant was discarded, and the mitochondria pellet was washed in liver isolation solution twice with centrifugation at 3,500 *g* for 10 minutes. The final mitochondria pellet was suspended in a mannitol-sucrose solution.

Mitochondrial respiration was determined polarigraphically (Oxytherm, Hansatech Instruments, UK) following procedures outlined previously [31]. In one chamber, respiration was measured using 2 mM pyruvate, 2 mM malate, and 10 mM glutamate as a substrate. In the second chamber, respiration was measured using 5 mM succinate as a substrate. State 2 respiration was defined as the respiration rate in the presence of substrates, and state 3 respiration, a measure of maximal respiration, was defined as the rate of respiration following the addition of 0.25 mM ADP to the chamber containing buffered mitochondria and respiratory substrates, and state 4 respiration was defined as the respiration rate measured after the phosphorylation of added ADP was complete. State 2, 3, and 4 respirations were measured at 40oC and were normalized to mitochondrial protein content. The respiratory control ratio (RCR) was calculated by dividing state 3 respiration by state 4 respiration.

Mitochondrial membrane potential was measured as described by [32]. Briefly, mitochondrial membrane potential was followed using the potential-sensitive dye safranin O [33]. Isolated mitochondria were incubated in standard buffer containing 3 mM HEPES, 1 mM EGTA, 0.3% (W/V) BSA, 1 µg/ml oligomycin, and 120 mM potassium chloride (pH = 7.2 and 40°C). Mitochondria were incubated at a concentration of 0.35 mg/ml mitochondrial protein in standard buffer with 5 µM safranin O. The change in fluorescence were measured in cuvette by Spectramax M (Molecular Devices, Sunnyvale, CA) at an excitation of 533 nm and an emission of 576 nm. In the end of each run, membrane potential was dissipated by addition of 2 µM FCCP. The relative decrease in fluorescent signal on energization of the mitochondria is used to represent the membrane potential. Results are reported as the absolute magnitude of this change in fluorescence, with larger changes in relative fluorescence units indicating higher membrane potentials.

The measurement of H_2_O_2_ emission in isolated mitochondria was conducted using Amplex Red (Thermofisher, Waltham, MA) [31]. Formation of resorufin (Amplex Red oxidation) by H_2_O_2_ was measured at an excitation wavelength of 545 nm and an emission wavelength of 590 nm using a Synergy H1 Hybrid plate reader (BioTek; Winooski, VT, USA), at 40°C in a 96-well plate using succinate. To eliminate carboxylesterase interference, 100 µM of phenylmethyl sulfonyl fluoride were added into experimental medium immediately prior measurement according to Miwa et al [34]. Readings of resorufin formation were recorded every 5 minutes for 15 minutes, and a slope (rate of formation) was produced from these. The obtained slope was then converted into the rate of H_2_O_2_ production using a standard curve and were normalized to mitochondrial protein levels. Citrate synthase activities was measured in liver homogenate as a function of the increase in absorbance from 5,5′-dithiobis-2-nitrobenzoic acid reduction according to Trounce et al [29].

Western blots were conducted on liver samples to analyze a marker of lipid peroxidation (4-Hydroxynonenal; 4-HNE; ab46545; Abcam, Cambridge, MA), a marker of protein oxidation (protein carbonyls; OxyBlot; s7150; EMD Millipore, Billerica, MA), and a marker of mitochondrial biogenesis (PGC-1*α*, GTX37356; Genetex, Irvine, CA). Each membrane was stained by Ponceau S and was used as the loading and transfer control. A chemiluminescent system was used to visualize marked proteins (GE Healthcare Life Sciences, Pittsburgh, PA). Images were taken and analyzed with the ChemiDocIt Imaging System (UVP, LLC, Upland, CA).

### (f) Analyses of tissue carotenoid content

We used high performance liquid chromatography (HPLC) to examine the carotenoid content of mitochondrial fractions proposed to be inner mitochondrial membrane, outer mitochondrial membrane, and matrix. The pelleted fractions were resuspended in 0.5 ml mitochondrial extraction buffer, and we subsampled 50 µl of the resuspension and extracted with 250 µl of ethanol, 500 µl water, and 1.5 ml of hexane:tert-Butyl ethyl ether 1:1 vol:vol. We collected the organic phase from the samples, dried this under a stream of nitrogen gas, then resuspended in 200 µl of mobile phase. We injected 50 µl of the resuspended extract in to an Agilent 1100 series HPLC equipped with a YMC carotenoid 5.0 µm column (4.6 mm × 250 mm, YMC). We eluted the samples with a gradient mobile phase consisting of acetonitrile:methanol:dichloromethane (44:44:12) (vol:vol:vol) through 11 minutes then a ramp up to acetonitrile:methanol:dichloromethane (35:35:30) from 11-21 minutes followed by isocratic conditions through 35 minutes. The column was held at 30°C, and the flow rate was 1.2 ml/min throughout the run. We monitored the samples with a photodiode array detector at 400, 445, and 480 nm, and carotenoids were identified by comparison to authentic standards or published accounts. Carotenoid concentrations were determined based on standard curves established with astaxanthin (for ketocarotenoids) and zeaxanthin (for xanthophylls) standards.

### (g) Statistical analyses

Statistical analyses were completed with SigmaStat 3.5, Systat Software, Inc., Point Richmond, CA, USA (fraction data), SAS 9.4, SAS Institute Inc, Cary, NC, USA, and R version 3.3.2, Vienna, Austria [35]. Analysis of variance was used to compare the relative concentration of mitochondrial membrane and matrix markers, and ketolated carotenoids between mitochondrial fractions. To compare the hue of the house finches to mitochondrial variables, we used a multivariate statistical model because the mitochondrial performance variables are dependent on one another. We used backward stepwise selection procedure for multiple regression to evaluate this relationship. For all analyses, significance was established at *p* < 0.05. The stepwise model was run in SAS and the model estimated coefficients from our final model were double-checked in R. In addition, we ran linear regression between variables in SAS to confirm biologically relevant relationships.

### (h) CYP2J19 molecular modeling

The amino acid sequence from House Finch CYP2J19 was submitted to the Robetta and I-TASSER servers for prediction of the protein’s 3D structure based on a combined *ab initio* and homology modeling approach [36,37]. The highest ranked output model from each server was utilized for analysis and cross examined for consistency with the other models.

The position of the predicted heme functional group of CYP2J19 was modeled through superimposition with the structure of the nearest structurally characterized homologue (CYP2B4, PDB ID = 3TK3). The position of this heme group is consistent with coordination by cysteine 444 of CYP2J19. Potential substrate tunnels were identified using Caver 3.0 [38], with the Fe atom of the heme functional group set as the starting point for the search starting point. A minimum probe radius of 0.9 Å, a shell depth of 4, a shell radius of 3 and a clustering depth of 3.5 was utilized. Identified tunnels were screened manually for likely involvement in substrate capture, based on size and proximity of the tunnel exit to the putative lipid embedded portion of CYP2J19. To validate the selected substrate tunnel, its position was compared to ligands in previous solved cytochrome P450 structures (PDB codes utilized: 4H1N, 3C6G, 4UFG, 3EBS, 1R90, 4I8V, 5X24, 4R20, 5T6Q, 3UA1, 2Q9F, 1ZOA, 4KEY, 2UWH and 4KPA). P450s bound to compact ligands accommodated them in a distinct region below the heme ligand, while elongated substrates clustered on the opposite side. The substrate tunnel predicted in CYP2J19 corresponded to the position of these larger substrates, which is consistent with the elongated nature of the carotenoid substrates.

In order to determine the likelihood that the N-terminus of CYP2J19 forms a lipid anchor, we analyzed the primary amino acid sequence and found that, typical of lipid anchors and secretion signal peptides, the first 45 amino acids are composed of predominantly hydrophobic residues. Both modelling programs, as well as the secondary structure prediction program JPRED, predicted that the majority of this region is α-helix in structure. This suggests that the N-terminus of CYP2J19 indeed forms a lipid anchoring helix. The models and associated analysis are contained in supplemental files associated with the manuscript.

## Results and discussion

### (a) Red carotenoids are localized in the inner mitochondrial membrane in molting house finches

A key prediction of the hypothesis that carotenoid coloration is linked to cellular respiration is that red carotenoid pigments should be present within mitochondria [19,25]. We previously documented that high levels of red ketolated carotenoids are associated with mitochondria in the livers of molting house finches [39], but we could not rule out that these pigments were actually concentrated in external mitochondrial-associated membranes. To resolve this, we isolated and fractionated hepatic mitochondria into IMM, OMM, and matrix, and subsequently measured the concentrations of the red carotenoid 3HE in the fractions.

Protein markers confirmed the separation of mitochondrial components (figure 3C). Consistent with our hypothesis, 3HE was present in high concentrations in the IMM (2.19 µg mL-1) (figure 3A and 3B). These levels were significantly higher than in the matrix (by 22-fold; *p* = 0.009) and OMM (by 7.8-fold; *p* = 0.014) fractions. The high levels of 3HE in the IMM fraction of mitochondria is strong evidence that ketolated carotenoids are not merely associated with mitochondria, but instead abundant in the interior membrane. This observation is of potentially great significance for understanding the mechanisms that underlie honest signaling via red carotenoid coloration. Given carotenoids compounds are co-localized with the electron transport system in the IMM, they are ideally localized within the cells of house finches to signal respiratory performance during sexual displays. These findings contrast with mammals, which do not use ketolated carotenoids in social signaling [42]; most mammals use enzymatic mechanisms to specifically exclude carotenoids from the mitochondria, including carotenoid-cleaving enzyme β,β-carotene-9,10-dioxygenase (BCO2) [41], and murine knockdowns of this gene have negative consequences for physiological function [40].

**Figure 3.**
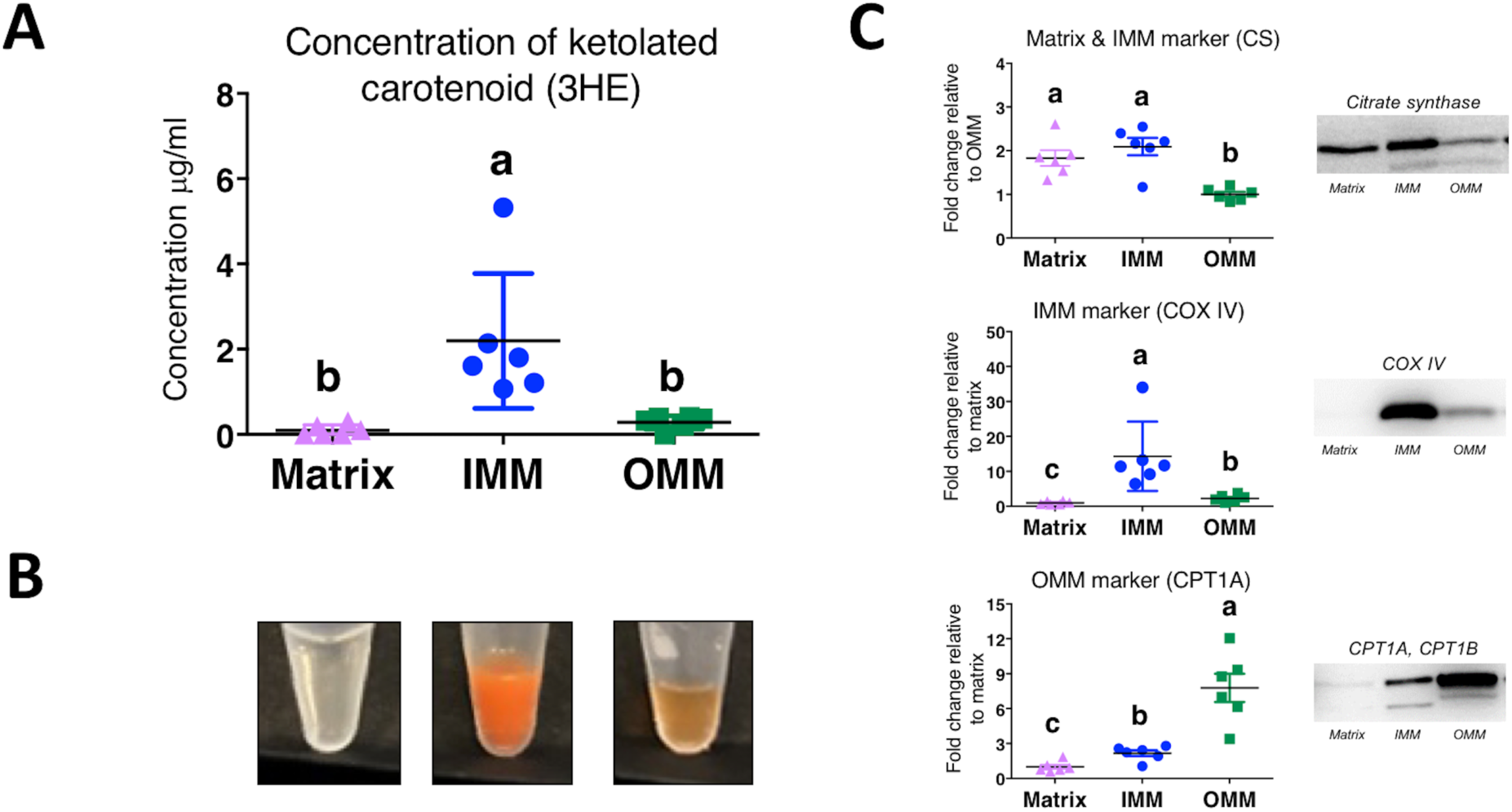
Ketolated carotenoid content of mitochondrial fractions from the livers of molting house finches. **(A)** The concentration of the red carotenoid 3-hydroxyechinenone (3HE) in each of three fractions of mitochondria isolated from house finch livers. 3HE concentration is higher in the inner mitochondrial membrane (IMM) than the matrix or outer mitochondrial membrane (OMM). **(B)** The relative 3HE content of each fraction was also evident in the color of fractions. **(C)** The separation of the three mitochondrial fractions was verified by western blotting of citrate synthase (CS), a citric acid enzyme located primarily in the matrix that also binds to the IMM, cytochrome *c* oxidase (COX IV), a protein of the electron transport system in IMM, and carnitine palmitoyltransferase I (CPT1), an enzyme that is unique to the OMM. Different lower-case letters above the data indicate significant differences among treatment groups (*p* < 0.05). Standard deviation bars are given. Western blots from which protein concentrations were estimated are given.

### (b) Carotenoid coloration is strongly correlated with mitochondrial function in house finches

The second key prediction of our central hypothesis is that bird coloration is positively associated with mitochondrial bioenergetic capacity [9,25]. We tested this prediction by capturing wild house finches that were actively growing their red feathers, quantifying the coloration of growing feathers, and measuring the performance of functional hepatic mitochondria. We measured isolated liver mitochondrial state 2 (proton leak), 3, and 4 respiration rates, RCR, and mitochondrial membrane potential both in the presence of OXPHOS complex I substrates (pyruvate, malate, and glutamate) and OXPHOS complex II substrate (succinate). We also quantified isolated liver mitochondrial H_2_O_2_ production and whole liver tissue adduct levels of 4-hydroxynonenal (4-HNE; a by-product of lipid peroxidation), protein carbonyls, citrate synthase activity, and PGC-1*α* protein levels (the mitochondrial biogenesis transcriptional activator).

We ran a stepwise-backwards regression model to determine which of the 15 variables made a significant contribution to the coloration of growing feathers. Six variables contributed to the best fit statistical model (F = 10.6, df = 6,25, *p* < 0.001, R2=0.727); RCR, state 2 respiration, state 4 respiration, and mitochondrial membrane potential with complex I substrates; 4-HNE levels and PGC-1α protein levels (figure 4, table 1). The strongest associations in both the backward regression model and the full multiple regression model were between the redness of growing feathers and both RCR and PGC-1*α* (figure 4).

**Figure 4.**
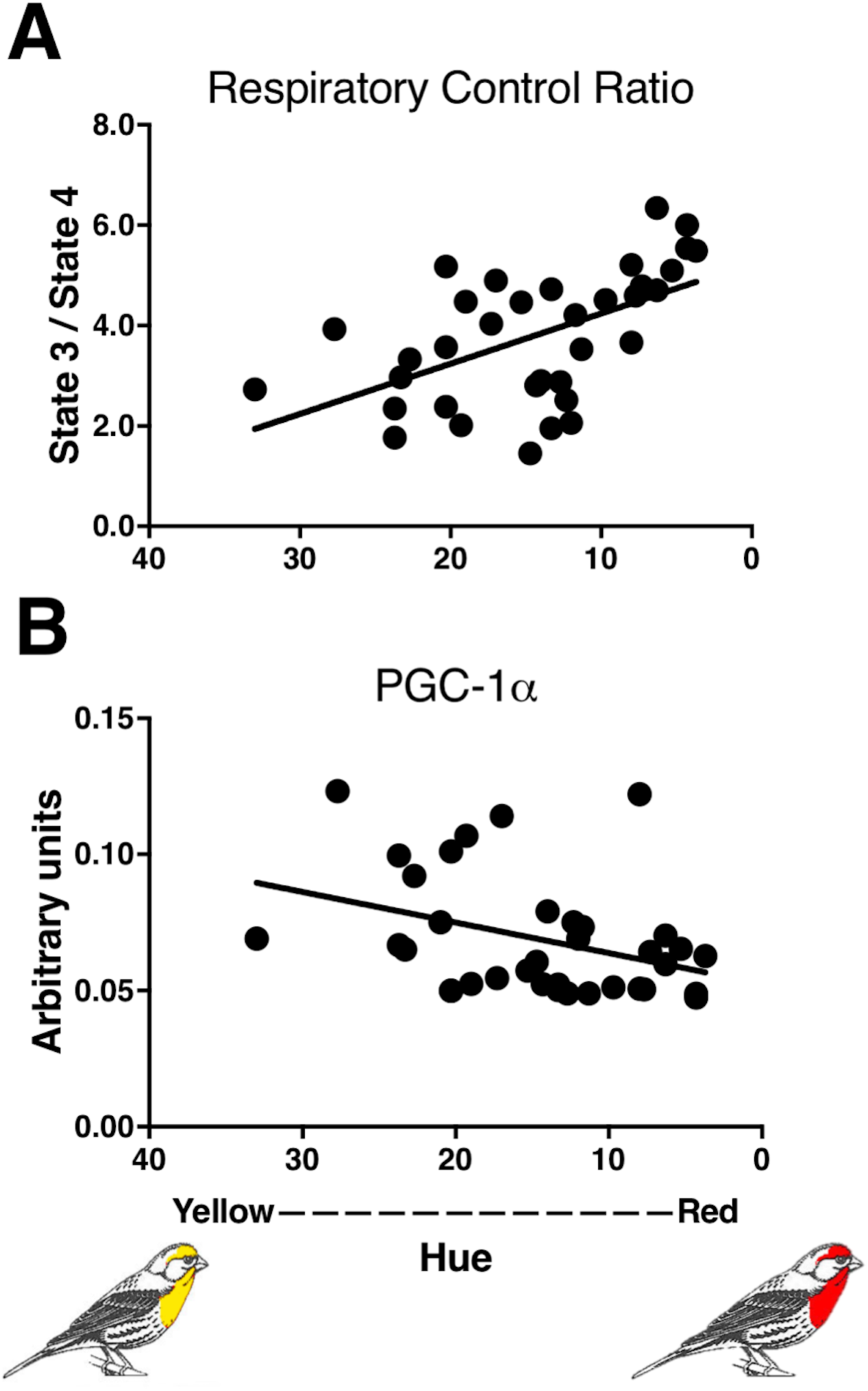
The relationships between feather redness and mitochondrial energy capacity and biogenesis. The results of a backward stepwise regression analysis (F = 10.6, df = 6,25, *p* < 0.001, R2=0.727) suggest that redness was most strongly correlated with (**A**) respiratory control ratio (P< 0.001) and (**B**) PGC-1*α* protein levels (P<0.001). Results of the full model and backwards regression are given in table 1.

The ten reddest birds had an average RCR 1.7-fold greater than the ten yellowest individuals (*p* < 0.001). The RCR is a measure of mitochondrial coupling efficiency; low ratios are associated not only with reduced performance but also with disease and aging in birds and mammals [45,46]. RCR is a ratio of state 3 divided by state 4 respiration, thus an increase in RCR should be associated with an increase in state 3 (maximum respiratory performance), a decrease in state 4 (basal, non-phosphating, respiratory performance), or both. In this study, RCR was greater in redder birds and, counter-intuitively, state 4 respiration was also higher in redder birds than dull birds in the backward-regression, but not full model. Given the inconsistency of the full and backward regression models, we interpret the state 4 values relative to RCR with caution. A direct comparison of the linear relationship between RCR and both state 3 and state 4 respiration is more revealing and clearly suggests that higher RCR is associated with lower state 4 respiration (linear regression, F_31_=45.8, *p*<0.001) but not higher state 3 respiration (linear regression, F_31_=0.39, *p*=0.53). This indicates that the hepatic mitochondria of redder birds appear to have lower costs associated with supporting basal respiratory than duller birds. Further, red birds also had increased mitochondrial membrane potential with complex I in both regression models and less leak (state 2 respiration) in the backward regression model and a near significant trend in the full model (table 1). These findings are also consistent with higher coupling efficiency associated with red coloration and supports the prediction that red birds have better mitochondrial function than drab birds.

We found a strong negative association between PGC-1*α* and plumage redness (figure 4B) that was consistent across the backward and full multiple regression models. PGC-1*α* is thought to be the master regulator of mitochondrial biogenesis, but it has also been associated with mitochondrial remodeling [47]. While both red and dull birds maintained comparable mitochondrial density (as indicated by consistent citrate synthase activity), higher PGC-1*α* suggests that the mitochondria of dull birds require more frequent replacement. Redder males also had significantly higher 4-HNE levels indicating that they produce more oxidants than drabber males (table 1). A modest increase in oxidant levels has been shown to increase relative antioxidant levels, upregulate repair mechanisms, and upregulate mitochondrial biogenesis, while high levels of oxidant typically lead to persistent oxidative damage [31,48–51]. Despite the fact that 4-HNE and PGC-1*α* had opposing relationships with hue, they had a positive linear relationship with each other (linear regression, F_31_=5.77, *p*=0.023) indicating that greater oxidative damage stimulates greater mitochondrial biogenesis, which would compensate for damaged mitochondria or replace mitochondria lost to mitophagy and apoptosis. Whether higher 4-HNE reflects a persistent detrimental response in these birds is unknown. Prior studies have suggested that in the short-lived house finch, redder birds have improved overwinter survival and improved ability to combat disease relative to dull birds [52,53].

### (c) An enzymatic model for how carotenoid coloration may be controlled by mitochondrial bioenergetics

Collectively, our observations indicate that carotenoid coloration is strongly correlated with hepatic mitochondrial performance in the house finch. In turn, it is plausible that respiratory performance may control carotenoid ketolation through processes occurring in the IMM, where 3EH is localized. One way in which these processes may be linked is through the putative cytochrome P450 enzyme CYP2J19, which has been shown to genetically required for the production of red carotenoid pigments in birds and turtles [21,22,57]. Based on this genetic link we hypothesized that this protein is an enzyme that catalyzes ketolation of yellow dietary carotenoids to their red derivatives in a reaction requiring an electron donor (e.g. NAD(P)H) and oxygen [21] (figure 2). This requirement for reducing equivalents produced in the mitochondria for carotenoid conversion could serve as a link between mitochondrial function and pigmentation.

As an initial exploration of this hypothesis and to demonstrate the plausibility of a mechanistic link between the ketolation of carotenoid pigments and aerobic respiration in cells, we used an *in silico* approach to create a molecular model of CYP2J19 (figure 5A, movie S1). This model strongly suggests that, like many other cytochrome P450s, CYP2J19 possesses an N-terminal anchor tethering it to a cellular membrane (figure 5A). As carotenoids are lipophilic, this localization would position CYP2J19 ideally for performing the ketolation reaction. We identified a substrate binding channel in CYP2J19 leading to its heme cofactor (figure 5A,C). This tunnel emerges from the region of CYP2J19 predicted to be embedded in the lipid bilayer, based on analysis of membrane-associated cytochrome P450 enzymes [58,59]. It is lined by largely hydrophobic residues and is of dimensions compatible with carotenoid binding (figure 5B,C). Each of these structural characteristics of CYP2J19 support the hypothesis that it catalyzes the conversion of yellow dietary carotenoids to red ketolated carotenoids in birds.

**Figure 5.**
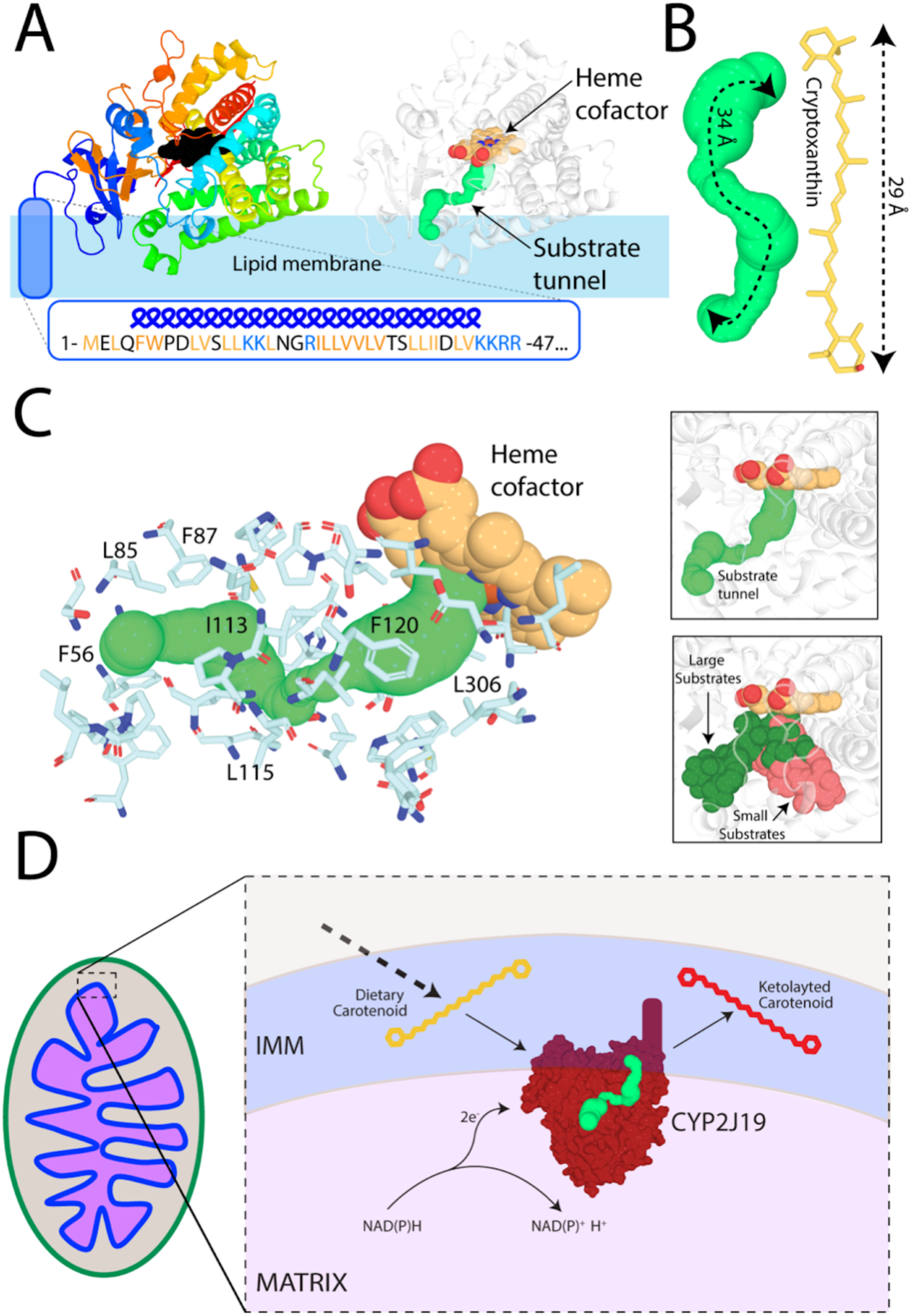
A possible molecular mechanism for the ketolation of carotenoids in the inner mitochondrial membrane. **(A)** A molecular model of the cytochrome P450 monooxygenase CYP2J19. Left: Cartoon ribbons colored from blue (N-terminal) to red (C-terminal), heme cofactor in black, and predicted N-terminal lipid anchor in blue. Right: Same model colored white with substrate tunnel shown as green spheres (right). **(B)** The length of the CYP2J19 substrate binding tunnel (left) is comparable to that of the putative carotenoid substrate cryptoxanthin (right). (**C**) The predicted substrate tunnel in CYP2J19 is surrounded by predominantly hydrophobic amino acids sidechains. Side chains surrounding tunnel are shown as sticks with representative hydrophobic residues labelled. The predicted CYP2J19 substrate tunnel (left) corresponds to the position of large substrates in previously solved cytochrome P450 structures (right). **(D)** A model for how the putative ketolation reaction performed by CYP2J19 may link red coloration to respiratory performance.

In turn, these characteristics of CYP2J19 support the hypothesis that its activity could directly link mitochondrial performance with carotenoid ketolation (figure 5D). Several scenarios are possible. At present, it is unknown whether or not CYP2J19 localizes within mitochondria, but if CYP2J19 is anchored to the IMM, then its catalytic function would be affected by various aspects of mitochondrial performance; the enzyme would be particularly sensitive to mitochondrial redox state and would share an electron carrier with other key mitochondrial enzymes if it is NADH-dependent (i.e. complex I) or NADPH-dependent (e.g. thioredoxin or glutathione reductases) (figure 5D). Alternatively, yellow carotenoids may be oxidized by CYP2J19 in the ER before being imported into the IMM or, as suggested previously [25], may be modified due to respiratory activity or ROS production independently of CYP2J19. Future experiments should focus on determining the localization of CYP2J19, the biochemical role it plays in carotenoid ketolation, and how its activity responds to aerobic respiration.

## 4. Conclusions

In this work, we demonstrate that (i) red carotenoids are localized in the inner mitochondrial membrane and (ii) plumage coloration is strongly correlated with mitochondrial function. Thus, we conclude that plumage coloration signals mitochondrial function, and hence core cellular functionality, in the house finch. Moreover, we propose the hypothesis that mitochondrial activity directly controls carotenoid ketolation. Detailed biochemical studies are needed to further test this latter hypothesis and it remains plausible that mitochondrial activity indirectly rather than directly controls carotenoid oxidation. The strength of the correlations between plumage coloration and mitochondrial function, the co-localization of the red carotenoids and the mitochondrial respiratory chain, and the molecular models predicting membrane localization and carotenoid compatibility of CYP2J19 all support a functional link between aerobic respiration in mitochondria and production of red ornamental coloration.

Linking ornamental feather coloration with mitochondrial function suggests a possible solution to a long-standing puzzle in evolutionary and behavioral biology: *what maintains the honesty of signals of individual condition?* [13]. Carotenoid coloration has been documented to signal a wide range of measures of individual performance, such as foraging ability, overwinter survival, immune system function, predator avoidance, and cognition [3,5]. In turn, mitochondrial function is a critical component of these same processes. Linking the ornamentation used in mate choice to function of core respiratory processes provides a novel mechanistic explanation for why carotenoid coloration relates to a range of aspects of individual performance and why females use plumage redness as a key criterion in choosing mates.

## Authors contributions

G.E.H. contributed to conceptualization, visualization, supervision, writing, and editing. W.R.H. contributed to conceptualization, visualization, supervision, funding acquisition, writing, and editing. J.D.J. contributed to conceptualization and editing. Z.G., N.R.P., H.A.T., V.A.A., M.J.P., N.M.J, and H.A.P. contributed to investigation and editing. R.G. and C.G. contributed to methodology, investigation, visualization, writing, and editing. A.N.K. contributed to methodology, supervision, writing, and editing. Y.Z. contributed to conceptualization, methodology, investigation, data curation, visualization, formal analysis, writing, and editing.

## Data accessibility

All data are available from the Dryad database (will be submitted after manuscript is accepted, accession number TBA).

## Acknowledgments

Maxwell Williams and Jordan Marquez assisted with trapping and laboratory assays. Matthew Toomey and Joseph Corbo conducted carotenoid analysis of mitochondrial fractions. Ryan Weaver, Chloe Josefson, and Kyle Heine provide valuable discussion and feedback. Vincent Careau suggested how to present the results of the backward regression model.

